# Screening for novel RyR2 inhibitors by ER Ca^2+^ monitoring

**DOI:** 10.1101/2023.08.21.553510

**Authors:** Mai Takenaka, Masami Kodama, Takashi Murayama, Mari Ishigami-Yuasa, Shuichi Mori, Ryosuke Ishida, Junji Suzuki, Kazunori Kanemaru, Masami Sugihara, Masamitsu Iino, Aya Miura, Hajime Nishio, Sachio Morimoto, Hiroyuki Kagechika, Takashi Sakurai, Nagomi Kurebayashi

## Abstract

Type 2 ryanodine receptor (RyR2) is a Ca^2+^ release channel on the endoplasmic/sarcoplasmic reticulum (ER/SR) that plays a central role in the excitation-contraction coupling in the heart. Hyperactivity of RyR2 has been linked to ventricular arrhythmias in patients with catecholaminergic polymorphic ventricular tachycardia (CPVT) and heart failure, where spontaneous Ca^2+^ release via hyperactivated RyR2 depolarizes diastolic membrane potential to induce triggered activity. In such cases, drugs that suppress RyR2 activity are expected to prevent the arrhythmias, but there is no clinically available RyR2 inhibitors at present. In this study, we searched for RyR2 inhibitors from a well-characterized compound library using a recently developed ER Ca^2+^-based assay, where the inhibition of RyR2 activity was detected by the increase in ER Ca^2+^ signals from R-CEPIA1er, a genetically encoded ER Ca^2+^ indicator, in RyR2-expressing HEK293 cells. By screening 1535 compounds in the library, we identified three compounds (chloroxylenol, methyl orsellinate and riluzole) that greatly increased the ER Ca^2+^ signal. All of the three compounds suppressed spontaneous Ca^2+^ oscillations in RyR2-expressing HEK293 cells and correspondingly reduced the Ca^2+^-dependent [^3^H]ryanodine binding activity. In cardiomyocytes from RyR2-mutant mice, the three compounds effectively suppressed abnormal Ca^2+^ waves without substantial effects on the action-potential-induced Ca^2+^ transients. These results confirm that ER Ca^2+^-based screening is useful for identifying modulators of ER Ca^2+^ release channels and suggest that RyR2 inhibitors have potential to be developed as a new category of antiarrhythmic drugs.

**Significance statement:** We successfully identified three compounds having RyR2 inhibitory action from a well-characterized compound library using an ER Ca^2+^-based assay, and demonstrated that these compounds suppressed arrhythmogenic Ca^2+^ wave generation without substantially affecting physiological action-potential induced Ca^2+^ transients in cardiomyocytes. This study will facilitate the development of RyR2 specific inhibitors as a potential new class of drugs for life-threatening arrhythmias induced by hyperactivation of RyR2.

## Introduction

Ryanodine receptors (RyRs) are Ca^2+^ release channels in the endoplasmic/sarcoplasmic reticulum (ER/SR) and three genetically distinct isoforms of RyR (RyR1–3) with 65-70% amino acid homology have been identified in mammalian tissues. Among them, RyR2 is indispensable in cardiac excitation-contraction coupling (Bers, 2001; Keefe et al., 2023; Woll and Van Petegem, 2022), where RyR2 is activated by Ca^2+^ entry through L-type Ca^2+^ channel (LTCC) via a Ca^2+^ induced Ca^2+^ release (CICR) mechanism. On the other hand, RyR1 plays a key role in skeletal muscle contraction and opens by depolarization-induced Ca^2+^ release mechanism via physical interactions with dihydropyridine receptor (DHPR) (Rios and Pizarro, 1991). RyR3 is present in low abundance in various tissues, and several physiological roles have been reported.

Genetic mutations in RyR2 are associated with various arrhythmogenic heart diseases, including catecholaminergic polymorphic ventricular tachycardia (CPVT), idiopathic ventricular tachycardia (IVF), and long QT syndrome (LQTS) (Hirose et al., 2022; Medeiros-Domingo et al., 2009; Nozaki et al., 2020; Priori and Chen, 2011; Sun et al., 2021; Tester et al., 2004; Woll and Van Petegem, 2022). Of these, CPVT mutations are of the gain-of-function type and account for approximately 90% of diseases linked to RyR2 mutations. When gain-of-function-type mutant RyR2 is further activated under strong sympathetic stimulation, spontaneous Ca^2+^ release occurs without Ca^2+^ entry from LTCC. This spontaneous Ca^2+^ release, in turn, activates inward sodium-calcium exchanger (NCX) currents to cause delayed afterdepolarization and triggered activity, leading to arrhythmias (Keefe et al., 2023; Lakatta, 1992; Tsien et al., 1979). In addition to the arrhythmic disorders linked to genetic mutations, it has been reported that excessive activation of RyR2 in chronic heart failure (HF) also causes arrhythmias (Benitah et al., 2021; Dridi et al., 2020; Szentandrassy et al., 2022).

The conventional antiarrhythmic drugs for ventricular arrhythmias in CPVT and HF include Na^+^ channel blockers, β-blockers and Ca^2+^ channel blockers. However, they sometimes fail to prevent sudden cardiac death. In arrhythmias induced by hyperactivation of RyR2, drugs that inhibit RyR channels are expected to effectively suppress the arrhythmias, but currently there are no clinically available RyR2 specific inhibitors. To date, several drugs, such as carvedilol (Zhou et al., 2011), flecainide (Watanabe et al., 2009), dantrolene (Kobayashi et al., 2010), EL20 (Klipp et al., 2018) and S107 (Lehnart et al., 2008), have been proposed to prevent the hyperactivity of mutated RyR2 (Szentandrassy et al., 2022). However, these drugs also interact with molecules other than RyR2. At present, it is difficult to predict how efficiently RyR2 inhibition itself can suppress arrhythmia. Thus, efforts to find RyR2 specific inhibitors are strongly required.

High-throughput screening (HTS) is a powerful method for rapidly evaluating many chemical compounds, which greatly accelerates drug discovery. We have reported that the ER Ca^2+^ concentration ([Ca^2+^]_ER_) in HEK293 cells expressing WT and mutant RyRs was inversely correlated with the Ca^2+^ release activity of RyRs at resting [Ca^2+^]_cyt_ (Kurebayashi et al., 2022; Murayama et al., 2016; Murayama et al., 2015), and developed an efficient high-throughput screening platform to find RyR1 inhibitors using HEK293 cells expressing mutant RyR1 and R-CEPIA1er (Murayama and Kurebayashi, 2019; Murayama et al., 2018).

In this study, we applied this method to identify RyR2 inhibitors by screening a chemical library of well-characterized drugs (1535 compounds). We successfully identified three compounds (chloroxylenol, methyl orsellinate, and riluzole) that prevent Ca^2+^ leakage from the ER to increase [Ca^2+^]_ER_ in HEK293 cells expressing RyR2. The three hit compounds exhibited RyR isoform specificities and dose-dependent RyR2 inhibition. These compounds suppressed Ca^2+^ waves in isolated cardiomyocytes without substantial effects on action-potential-induced Ca^2+^ transients. Our results indicate that RyR2 inhibitors are promising candidates for novel anti-arrhythmic drugs.

## Materials and Methods

### Generation of stable and inducible HEK293 cell lines expressing RyRs

HEK293 cells stably expressing ER Ca^2+^ sensor protein R-CEPIA1er (Suzuki et al., 2014) together with inducibly expressing wild-type (WT) RyR2 were generated as described previously (Kurebayashi et al., 2022; Murayama and Kurebayashi, 2019; Murayama et al., 2018). Briefly, a full-length mouse RyR2 cDNA was cloned in a tetracycline-induced expression vector (pcDNA5/FRT/TO; Life Technologies, Carlsbad, CA, USA) (Tong et al., 1997). Flp-In T-REx HEK293 cells (Thermo Fisher Scientific) were co-transfected with this expression vector and pOG44 Flp-recombinase expression vector in accordance with the manufacturer’s instructions. Clones with suitable doxycycline-induced expression of RyR2 were selected and used for experiments. The cells were then transfected with cDNA of R-CEPIA1er for its stable expression. Cells were cultured in Dulbecco’s Modified Eagle’s Medium supplemented with 10% fetal calf serum, 2 mM L-glutamine, 15 μg/ml blasticidin, 100 μg/ml hygromycin, and 400 μg/ml G418. When testing the effects of compounds on other RyR isoforms, we used HEK293 cells co-expressing RyR1 and R-CEPIA1er, and RyR3 and R-CEPIA1er that were generated by methods similar to those described above (Kurebayashi et al., 2022; Murayama and Kurebayashi, 2019; Murayama et al., 2018). To test the effects of compounds on mutant RyR2s, RyR2-R2474S, -R4497C, -R176Q, -N2386I, phospho-null triple mutant S2807A/S2813A/S2030A (RyR2-S3A) and phospho-mimetic triple mutant (RyR2-S3D) cells (Iyer et al., 2020; Kurebayashi et al., 2022; Uehara et al., 2017) were infected with baculovirus vector to express R-CEPIA1er.

### Time-lapse [Ca^2+^]_ER_ measurement using multi-well plate fluorometer

Time-lapse [Ca^2+^]_ER_ measurements were performed using the FlexStation 3 fluorometer (Molecular Devices, San Jose, CA, USA), as described previously (Murayama and Kurebayashi, 2019; Murayama et al., 2018). Briefly, HEK293 cells were seeded on 96-well, flat, clear-bottomed black microplates (#3603; Corning, New York, NY, USA) at day 1, expression of RyR2 was induced by adding doxycycline (2 μg/ml) to the culture medium on day 2, and time-lapse ER Ca^2+^ measurements were carried out on day 3. Before the measurements, the culture medium in individual wells was replaced with 90 μl of HEPES-Krebs solution (140 mM NaCl, 5 mM KCl, 2 mM CaCl_2_, 1 mM MgCl_2_, 11 mM glucose, and 5 mM HEPES, pH 7.4). R-CEPIA1er was excited at 560 nm and fluorescence emitted at 610 nm was captured every 10 s for 300 s. At 100 s after the start of measurement, 60 μl of compound solution, containing 25 μM test compound in HEPES-Krebs solution, was applied to the wells of the reading plate at a final concentration of 10 μM. The change in fluorescence induced by the compounds was expressed as *F/F_0_*, in which average fluorescence intensity of the last 100 s (*F*) was normalized to that of the initial 100 s (*F_0_*). We screened 1535 well-characterized drugs in the Tokyo Medical and Dental University (TMDU) chemical compound library at a concentration of 10 μM. Measurements were performed at 37.

### Analysis of high-throughput screening data

Assay quality was determined based on positive (1 mM tetracaine) and negative [0.1% dimethylsulfoxide (DMSO)] controls, as indexed by Z’ factor:

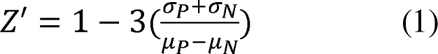

where *σ_P_* and *σ_N_* are the S.D.s of positive and negative controls, and *μ_P_* and *μ_N_* are the means of positive and negative controls, respectively.

### Reagents

The following reagents were purchased after finding hit compounds from the initial TMDU library screening. Chloroxylenol and methyl orserllinate (supply name: methyl 2,4-dihydroxy-6-methylbenzoate) were purchased from Toronto Research Chemicals (Toronto, Canada), riluzole (supply name: 2-amino-6-(trifluoromethyl)benzothiazole) and 4-chloro-3-ethylphenol (4-CEP) from Tokyo Chemical Industry (Tokyo, Japan), and 4-chloro-m-cresol (4-CMC, supply name: 4-chloro-3-methylphenol) from Fujifilm Wako Chemicals (Kyoto, Japan). All the above reagents were dissolved in dimethyl sulfoxide (DMSO) and used at a final concentration of 0.1% DMSO.

### Simultaneous measurement of [Ca^2+^]_ER_ and [Ca^2+^]_cyt_ in single HEK293 cells expressing RyR2s

For measurements of Ca^2+^ signals in individual HEK293 cells, [Ca^2+^]_cyt_ and [Ca^2+^]_ER_ were simultaneously monitored using genetically encoded Ca^2+^ indicators, G-GECO1.1 (Zhao et al., 2011) and R-CEPIA1er (Suzuki et al., 2014), respectively (Kurebayashi et al., 2022). Cells were transfected with G-GECO1.1 and R-CEPIA1er cDNA 26–28 h before measurements. Doxycycline was added to the medium at the same time as transfection. G-GECO 1.1 and R-CEPIA1er were excited at 488 nm and 561 nm, respectively, through a 20× objective lens and light emitted at 525 nm and 620 nm, respectively, was simultaneously captured with an EM-CCD camera at 700 ms intervals (Model 8509; Hamamatsu Photonics, Hamamatsu, Japan). Cytosolic ([Ca^2+^]_cyt_) and ER Ca^2+^ ([Ca^2+^]_ER_) signals were monitored for 4 min in HEPES-Krebs solution, 4 min in the presence of a test compound, 3 min in HEPES-Krebs solution, and then 1.5 min in 10 mM caffeine-containing Krebs solution. At the end of the measurement, the cells were perfused with the following solutions to obtain [Ca^2+^]_cyt_ and [Ca^2+^]_ER_ calibrations: 0Ca-Krebs solution (140 mM NaCl, 5 mM KCl, 1 mM MgCl_2_, 11 mM glucose, 10 mM HEPES, pH 7.4), BAPTA-0Ca-Krebs solution containing 5 mM 1,2-bis(o-aminophenoxy)ethane-N,N,N′,N′-tetraacetic acid (BAPTA) and 20 μM ionomycin, 0Ca-Krebs solution, and then 20Ca-Krebs solution containing 20 mM CaCl_2_ and 20 μM ionomycin (Kurebayashi et al., 2022). *F_min_* and *F*_max_ values were obtained with the BAPTA-0Ca-Krebs solution and 20Ca-Krebs solution, respectively.

To examine the effects of test compounds on [Ca^2+^]_cyt_ oscillations, cells were loaded with 4 µM fluo-4 AM in culture medium for 30 min at 37°C and then incubated with HEPES Krebs solution. Fluo-4 was excited at 488 nm through a 20× objective lens and light emitted at 525 nm was captured with the EM-CCD camera at 700 ms intervals. All measurements were performed at 26°C by perfusing solutions using an in-line solution heater/cooler (Warner Instruments, Holliston, MA, USA).

### [^3^H]Ryanodine binding

The assay was carried out as described previously (Kurebayashi et al., 2022). In brief, microsomes prepared from HEK293 cells stably expressing the RyR2s were incubated with 5 nM [^3^H]ryanodine for 1 h at 25 [in medium containing 0.17 M NaCl, 20 mM 3-morpholino-2-hydroxypropanesulfonic acid (MOPSO) (pH 7.0), 2 mM dithiothreitol, 1 mM adenosine monophosphate, 1 mM MgCl_2_, and various concentrations of free Ca^2+^ buffered with 10 mM EGTA. Free Ca^2+^ concentrations were calculated using WEBMAXC STANDARD (https://somapp.ucdmc.ucdavis.edu/pharmacology/bers/maxchelator/webmaxc/webmaxcS.htm). Protein-bound [^3^H]ryanodine was separated by filtration through polyethyleneimine-treated glass filters (Filtermat B; PerkinElmer, Waltham, MA) using a Micro 96 Cell Harvester (Skatron Instruments, Lier, Norway). Nonspecific binding was determined in the presence of 20 μM unlabeled ryanodine.

### Ca^2+^ imaging in isolated mouse single cardiac myocytes

All animal-handling procedures were in accordance with the guidelines and approved by the ethics committees of Juntendo University School of Medicine. For Ca^2+^ imaging in isolated cardiomyocytes, 2-month-old WT and homozygous RyR2-R420W (Okudaira et al., 2014) and 3-month-old homozygous *Tnnt2*-ΔK210 mice (Du et al., 2007; Odagiri et al., 2014) were used.

Mice were deeply anesthetized with pentobarbital sodium (100 mg/kg i.p.), and their hearts were excised and rinsed in Krebs solution. Single cardiac myocytes were isolated from the ventricles of mice using an established enzymatic method (Shioya, 2007) and loaded with Cal520-AM (AAT Bioquest Inc., Pleasanton, CA, USA). Cells were field-stimulated at 0.5 Hz in HEPES-buffered Tyrode’s solution (140 mM NaCl, 5 mM KCl, 1.5 mM CaCl_2_, 1 mM MgCl_2_, 0.3 mM NaH_2_PO_4_, 11 mM glucose, 10 mM HEPES, pH 7.4) with or without test compounds. Cal520 was excited at 488 nm through a 20× objective lens and light emitted at 525 nm was captured with an EM-CCD camera at 8–14 ms intervals (Model 8509; Hamamatsu Photonics). Fluorescence signals (*F*) of Cal520 in individual cells were determined using region of interest (ROI) analysis, from which the cell-free background fluorescence was subtracted. To confirm intracellular Ca^2+^ signals with high resolution, video images were acquired at 6 ms interval using a 60x objective lens. All measurements were carried out at 30°C by perfusing solutions through an in-line solution heater/cooler (Warner Instruments, Holliston, MA, USA).

### Statistics

Data are presented as the mean ± S.D. Statistical analysis was performed using Prism 9 (GraphPad Software, Inc., La Jolla, CA). Unpaired Student’s t test was used for comparisons between two groups. One-way analysis of variance (ANOVA) followed by Dunnett’s test was performed to compare multiple groups with one factor, while two-way ANOVA followed by Tukey’s test was used to compare multiple groups in experiments with two factors. Statistical significance was defined as P<0.05 when compared with the negative control. These analyses were performed as planned and recorded in the protocols.

## Results

### Validation of screening platform for RyR2 inhibitors

In this study, we used HEK293 cells stably expressing R-CEPIA1er (Suzuki et al., 2014) together with inducibly expressing WT RyR2 as a screening platform for RyR2 inhibitors. Figure 1A illustrates the concept of the ER Ca^2+^-based screening for RyR2 inhibitors. [Ca^2+^]_ER_ is generally determined by the balance between Ca^2+^ release via the Ca^2+^ release channels and Ca^2+^ uptake by sarcoplasmic/endoplasmic reticulum Ca^2+^-ATPase (SERCA) Ca^2+^ pumps (Kurebayashi et al., 2022; Murayama et al., 2018). We have demonstrated that expression of WT and CPVT mutant RyR2s reduce [Ca^2+^]_ER_ by spontaneous Ca^2+^ release and that [Ca^2+^]_ER_ is inversely correlated with the channel activity: channels with greater activity cause more reduced [Ca^2+^]_ER_ (Fujii et al., 2017; Kurebayashi et al., 2022). Because induction of RyR2 expression causes spontaneous Ca^2+^ release to moderately lower [Ca^2+^]_ER_ in HEK293 cells (Figs. 1A-a and 1A-b) (Fujii et al., 2017; Kurebayashi et al., 2022), the inhibition of RyR2 by drugs will increase [Ca^2+^]_ER_ by means of Ca^2+^ uptake (Fig. 1A-c). In contrast, further activation of RyR2 by drugs will enhance Ca^2+^ release, leading to a further decrease in [Ca^2+^]_ER_ (Fig. 1A-d).

**Fig. 1.**
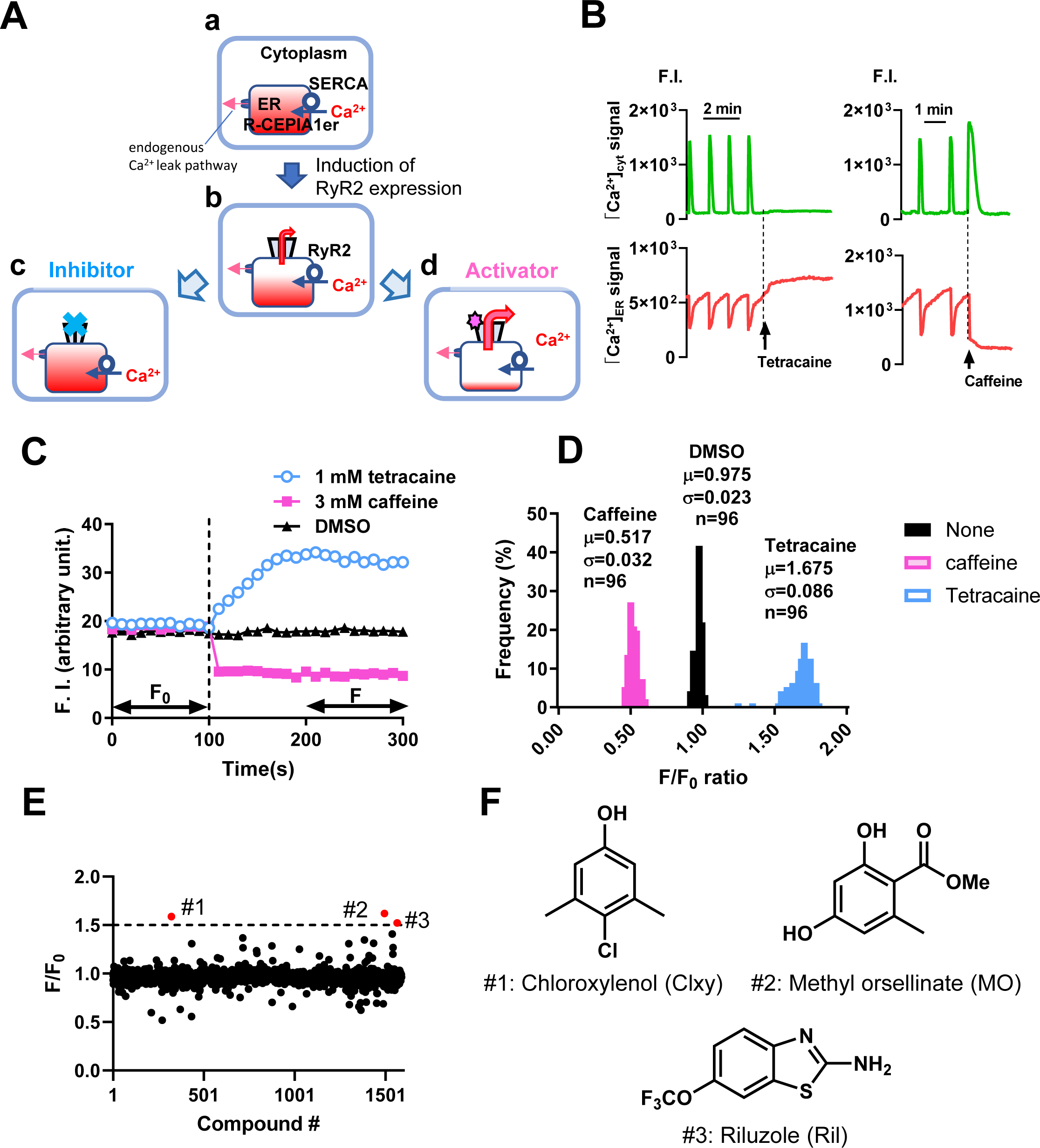
Screening of RyR2 inhibitors from well-characterized drug library by ER Ca^2+^ monitoring. (A) Scheme drawing of ER Ca^2+^-based detection of RyR2 activators and inhibitors. Induction of RyR2 moderately decreases [Ca^2+^]_ER_, from (a) to (b). RyR2 inhibitors and activators increase (c) and decrease (d) [Ca^2+^]_ER_, respectively. (B) Representative cytoplasmic (G-GECO1) and ER (R-CEPIA1er) Ca^2+^ signal changes upon application of tetracaine (left) and caffeine (right) in single HEK293 cells expressing WT RyR2. (C) Representative time-lapse R-CEPIA1er fluorescence measurement using a FlexStation3 fluorometer with HEK293 cells expressing WT RyR2. Test solutions containing DMSO (0.2%), caffeine (3 mM), and tetracaine (1 mM) were added at 100 s (dotted line) after the start of measurements. (D) Histograms of *F/F_0_* for DMSO, caffeine, and tetracaine (n = 96 wells each) in WT RyR2 cells. (E) Screening results. A TMDU chemical compound library of well-characterized drugs (1535 compounds, 10 μM) was screened in HEK293 cells expressing WT RyR2. Data are the mean of duplicate screens. The three compounds (#1-#3) with *F/F_0_* greater than 1.5 (dotted line) were selected as hits. (F) Chemical structures of the three hit compounds: chloroxylenol, methyl orsellinate, and riluzole.

Fig. 1B shows representative actual effects of tetracaine and caffeine, a known inhibitor and activator of ryanodine receptors (RyRs), respectively, on simultaneously measured [Ca^2+^]_cyt_ and [Ca^2+^]_ER_ signals in single HEK293 cells expressing WT RyR2. As previously reported, cells expressing RyR2 show spontaneous [Ca^2+^]_cyt_ oscillations and corresponding periodic [Ca^2+^]_ER_ decrease (Jiang et al., 2007; Kurebayashi et al., 2022; Uehara et al., 2017). The [Ca^2+^]_ER_ level was increased by an application of tetracaine (1mM) with termination of spontaneous [Ca^2+^]_cyt_ oscillations (left), whereas the [Ca^2+^]_ER_ was substantially decreased by the application of caffeine (3 mM) (right).

Figure 1C shows representative effects of tetracaine and caffeine on the R-CEPIA1er fluorescence signals in WT RyR2 cells measured with a FlexStation3 microplate reader. The cells were seeded in 96-well plates on the first day and RyR2 expression was induced by doxycycline on the next day. 24 h after the induction, R-CEPIA1er fluorescence signals were measured using the FlexStation3 fluorometer, which obtains the summation of fluorescence signals from many cells in the optical field (1mm diameter). The application of 3 mM caffeine rapidly decreased the R-CEPIA1er signal, whereas tetracaine (1 mM) gradually increased the fluorescence signal, which reached a plateau at around 200 s. The fluorescence ratio (*F/F_0_*) before and after the application of compounds was determined by normalizing the average fluorescence intensity for the last 100 s (*F*) to that for the initial 100 s (*F_0_*) (Fig. 1C). These results are consistent with the above idea that [Ca^2+^]_ER_ monitors RyR2 activity.

To quantitatively validate the assay system, we determined the coefficient of variation (CV), the ratio of the standard deviation to the mean and Z’ factor using 96 wells in each condition. Histograms of *F/F_0_* showed that fluorescence signals changed by tetracaine (*μ*=1.675 and *σ*=0.086) and caffeine (*μ*=0.517, *σ*=0.032), where *<u>μ</u>* and *σ* are the mean and S.D., respectively. were perfectly separated from that changed by DMSO (*μ*=0.975 and *σ*=0.023) (Fig. 1D). CV values for tetracaine, caffeine, and DMSO were 0.05, 0.06, and 0.02, respectively, which satisfied the criterion for HTS (CV<0.1). The Z’ factors were calculated by eq. 1 as 0.533 and 0.640 for tetracaine and caffeine, respectively (see Materials and Methods), which were within the acceptable range (Z’ > 0.5).

### Screening of a chemical compound library for RyR2 inhibitors by [Ca^2+^]_ER_ measurements

Using this screening platform based on [Ca^2+^]_ER_ measurements, we screened a TMDU chemical compound library of well-characterized drugs (1535 compounds) at 10 μM, which was the same as that used for the search for RyR1 inhibitors (Murayama et al., 2018). The overall results with WT RyR2 cells in duplicate assays are shown in Fig. 1E. We identified three compounds (#1–#3) with *F/F_0_* values greater than 1.5 (Fig. 1E). They were chloroxylenol (#1), methyl orsellinate (#2), and riluzole (#3) (Fig. 1F).

### Dose-dependent effects of hit compounds on WT and mutant RyR2s

To further characterize the hit compounds, we examined dose-dependent effects on [Ca^2+^]_ER_ in RyR2 WT cells using the same assay system (Fig. 2A). Chloroxylenol showed the highest potency (EC_50_ = 0.3 μM), followed by methyl orsellinate (EC_50_ = 1.1 μM) and riluzole (EC_50_ = 9.9 μM). Since RyR2 inhibitors are expected to be used for patients carrying mutant RyR2, we also tested these compounds on CPVT-linked mutant RyR2s, R2474S and R4497C, which strongly and moderately activate the channel activity, respectively (Kurebayashi et al., 2022). The three compound also similarly suppressed the CPVT-linked mutant RyR2s, R2474S and R4497C (Fig. 2B and C), as well as WT RyR2. The inhibitory effects of these compounds were also confirmed with arrhythmogenic right ventricular dysplasia (ARVD)/cardiomyopathy-associated mutant RyR2s such as R176Q (Kannankeril et al., 2006) and N2386I (Tiso et al., 2001), as well as RyR2 mutants at the three important phosphorylation sites (Huke and Bers, 2008; Lanner et al., 2010; Potenza et al., 2019), a phospho-null triple mutant S2807A/S2813A/S2030A (RYR2 S3A), and a phospho-mimetic triple mutant of RyR2 S2807D/S2813D/S2030D (RyR2 S3D) (Supplementary Fig. 1).

**Fig. 2.**
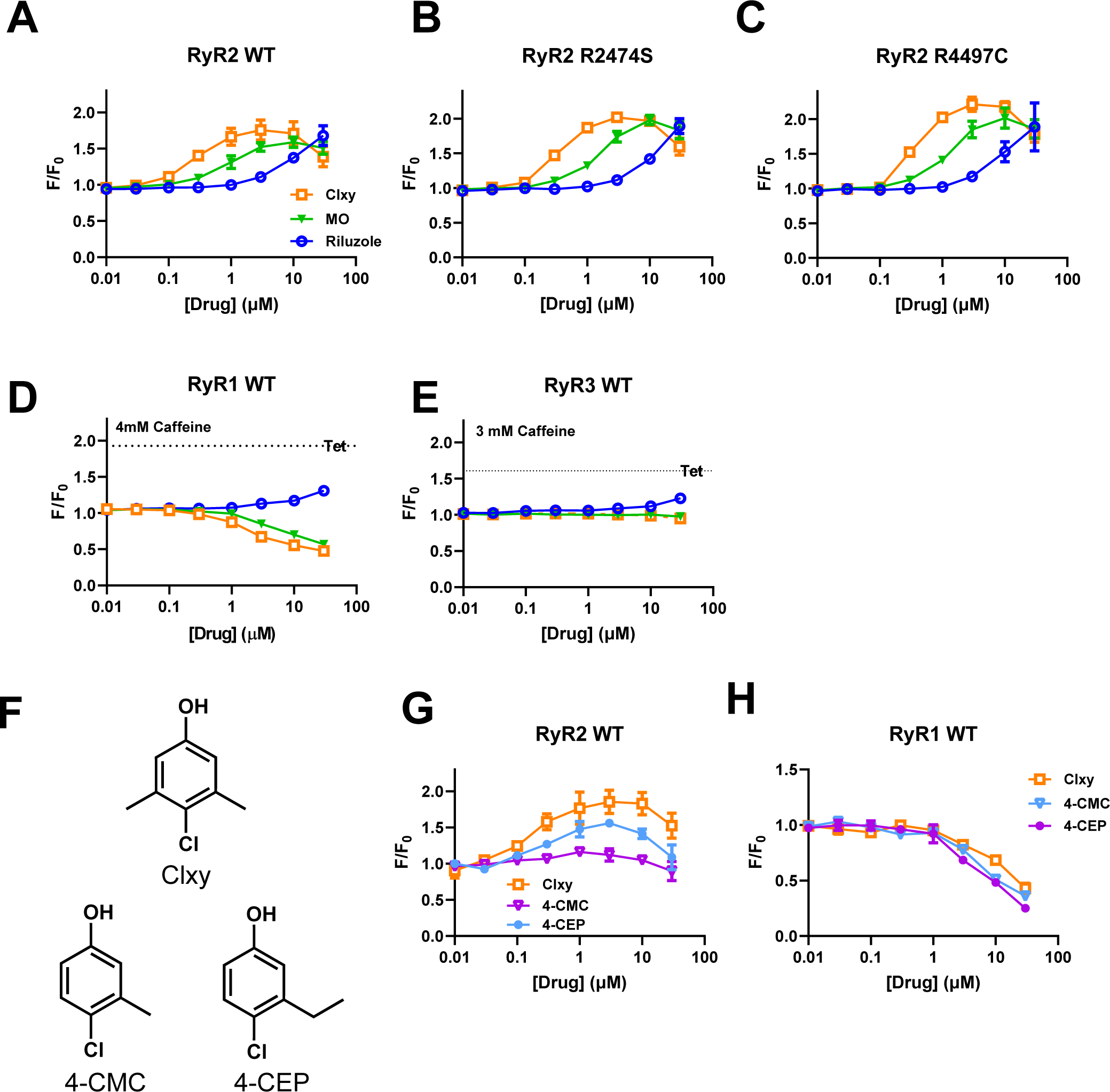
Dose-dependent effects of hit compounds on R-CEPIA1er signals in RyR-expressing cells determined with FlexStation3. (A) RyR2-WT, (B) RyR2-R2474S, (C) RyR2-R4497C, (D) RyR1-WT, (E) RyR3-WT. Measurements were carried out in normal Krebs solution for A–C, and in normal Krebs solution containing 4 mM and 3 mM caffeine for (D) and (E), respectively. (F) Chemical structures of chloroxylenol, 4-chloro-m-cresol (4-CMC) and 4-chloro-3-ethylphenol (4-CEP). (G) and (H) Dose-dependent effects of Clxy, 4-CMC, and 4-CEP on R-CEPIA1er signals in RyR2-WT (G) and RyR1-WT expressing cells (H). Data are mean ± SD (n=3–5).

To examine the RyR isoform specificity, the effects of these compounds on WT RyR1-WT and RyR3-WT cells were also examined (Fig. 2D and E). Because [Ca^2+^]_ER_ of RyR1-WT and RyR3-WT cells was almost fully loaded due to least spontaneous Ca^2+^ release activity (Murayama et al., 2018), measurements were performed in the presence of 4 mM (for RyR1-WT) or 3 mM (for RyR3-WT) caffeine, which enhances RyR channel activity to moderately reduce [Ca^2+^]_ER_. The addition of tetracaine increased [Ca^2+^]_ER_ (see dotted lines in Fig. 2D and E). Riluzole slightly increased [Ca^2+^]_ER_ in the RyR1-WT cells at 30 μM, whereas chloroxylenol and methyl orsellinate clearly reduced [Ca^2+^]_ER_ in the RyR1-WT at 3 μM or higher (Fig. 2D). These compounds had little effect on RyR3 (Fig. 2E). Therefore, the three compounds all suppress RyR2, although chloroxylenol and methyl orsellinate have some activating effects on RyR1.

The chemical structure of chloroxylenol is similar to those of 4-chloro-m-cresol (4-CMC) and 4-chloro-3-ethylphenol (4-CEP), which are well-known potent RyR1 activators (Westerblad et al., 1998; Zorzato et al., 1993) (Fig. 2F). We therefore examined the effects of these drugs on RyR1- and RyR2-expressing cells. The 4-CEP at 0.3–10 μM increased *F/F_0_* in RyR2 cells, indicating that 4-CEP also has some RyR2-inhibiting effect (Fig. 2G), whereas 4-CMC did not exhibit RyR2 inhibition. All of these compounds showed trends of decreasing *F/F_0_* of RyR2 cells from the peak at 30 μM, suggesting that they have a Ca^2+^-releasing effect on RyR2 at higher concentrations, i.e., 100∼300 μM or more. On RyR1-expressing cells, these compounds all similarly reduced *F/F_0_*, at 3 µM or more, indicating their Ca^2+^-releasing effects on RyR1 (Fig. 2H). These results indicate that the activating effect of chloroxylenol on RyR1 is similar to those of 4-CMC and 4-CEP (Westerblad et al., 1998; Zorzato et al., 1993).

### Effects of hit compounds on [^3^H]ryanodine binding

Next, we examined the effects of the three hit compounds on RyR2 activity via a Ca^2+^-dependent [^3^H]ryanodine binding assay (Fig. 3A). Since ryanodine specifically binds open RyR channels, [^3^H]ryanodine binding reflects the RyR2 channel activity (Fujii et al., 2017; Kurebayashi et al., 2022). Chloroxylenol (3 μM) and methyl orsellinate (10 μM) suppressed maximal activity and shifted the Ca^2+^ dependence to the right (Fig. 3A, left). Riluzole (10 and 100 μM) showed similar effects on [^3^H]ryanodine binding (Fig. 3A, right). Figure 3B shows the activity measured at pCa 5, a Ca^2+^ concentration close to that under physiological conditions. All three compounds suppressed [^3^H]ryanodine binding with IC_50_s similar to those obtained with the [Ca^2+^]_ER_-based assay (chloroxylenol, 0.13 μM; methyl orsellinate, 1.6 μM: riluzole, 5.3 μM).

**Fig. 3.**
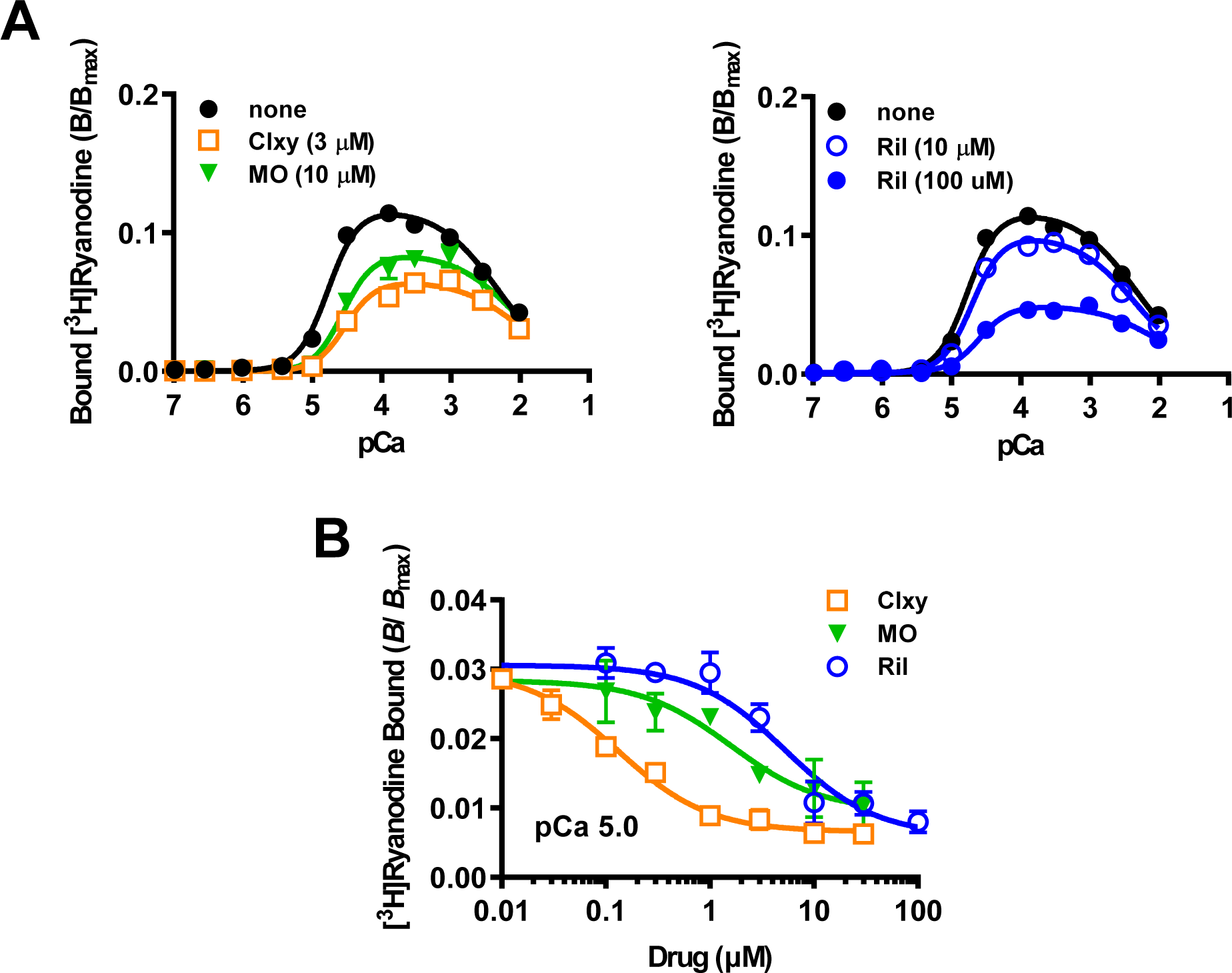
Effects of three hit compounds on [^3^H]ryanodine binding. (A) Effects of 3 μM chloroxylenol (Clxy, left), 10 μM methyl orsellinate (MO, left), and 10 and 100 μM riluzole (Ril, right) on Ca^2+^-dependent [^3^H]ryanodine binding. Data are mean ± SD (n=4–6). The curves are the results of fit of the equation to the data as follows,

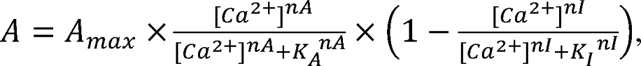

where *A* is the activity at the specified Ca^2+^, *A*_max_ is the gain that determines the maximal attainable activity, *K*_A_ and *K*_I_ are dissociation constants, and *n*_A_ and *n*_I_ are the Hill coefficients for Ca^2+^ of activation and inactivation, respectively. which are fixed at 2.0 and 1.0, respectively (Kurebayashi et al., 2022). The *A_max_*, *K_A_* and *K_I_* values obtained from the fitting are; none (0.118, 17 μM, 4.8 mM); 3 μM Clxy (0.066. 30 μM, 9.9 mM); 10μM MO (0.086, 28 μM, 8.3 mM); 10 μM Ril (0.106, 30 μM, 9.9 mM); 100 μM Ril (0.049, 25 μM, 9.1 mM). (B) Dose-dependent effects of chloroxylenol (Clxy), methyl orsellinate (MO), and riluzole (Ril) on [^3^H]ryanodine binding at pCa 5.0. Data are mean ± SD (n=2–4). The standard dose-dependent inhibition equation with the Hill slope of 1 were fitted to the data as follows,

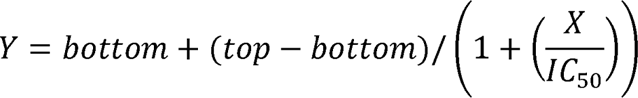 The IC_50_s from the fitting are Clxy (0.13 μM), MO (1.6 μM) and Ril (5.3 μM).

### Effects of hit compounds on single-cell Ca^2+^ homeostasis in HEK293 cells expressing RyR2

The above screening results indicate that RyR2 activity is suppressed by these hit compounds. Because HEK293 cells expressing RyR2 show spontaneous Ca^2+^ oscillations (Jiang et al., 2007; Kurebayashi et al., 2022; Uehara et al., 2017) as shown in Fig. 1B, we investigated how these compounds affect Ca^2+^ oscillations caused by RyR2. Figure 4A shows simultaneous measurements of [Ca^2+^]_cyt_ and [Ca^2+^]_ER_ signals from individual HEK293 cells expressing WT RyR2. Chloroxylenol (1 μM), methyl orsellinate (10 μM), and riluzole (10 μM) completely suppressed the spontaneous Ca^2+^ oscillations associated with the increased [Ca^2+^]_ER_ (Fig. 4B). The washout of the drugs immediately restored the Ca^2+^ oscillation, indicating that the effects of these compounds are reversible.

**Fig. 4.**
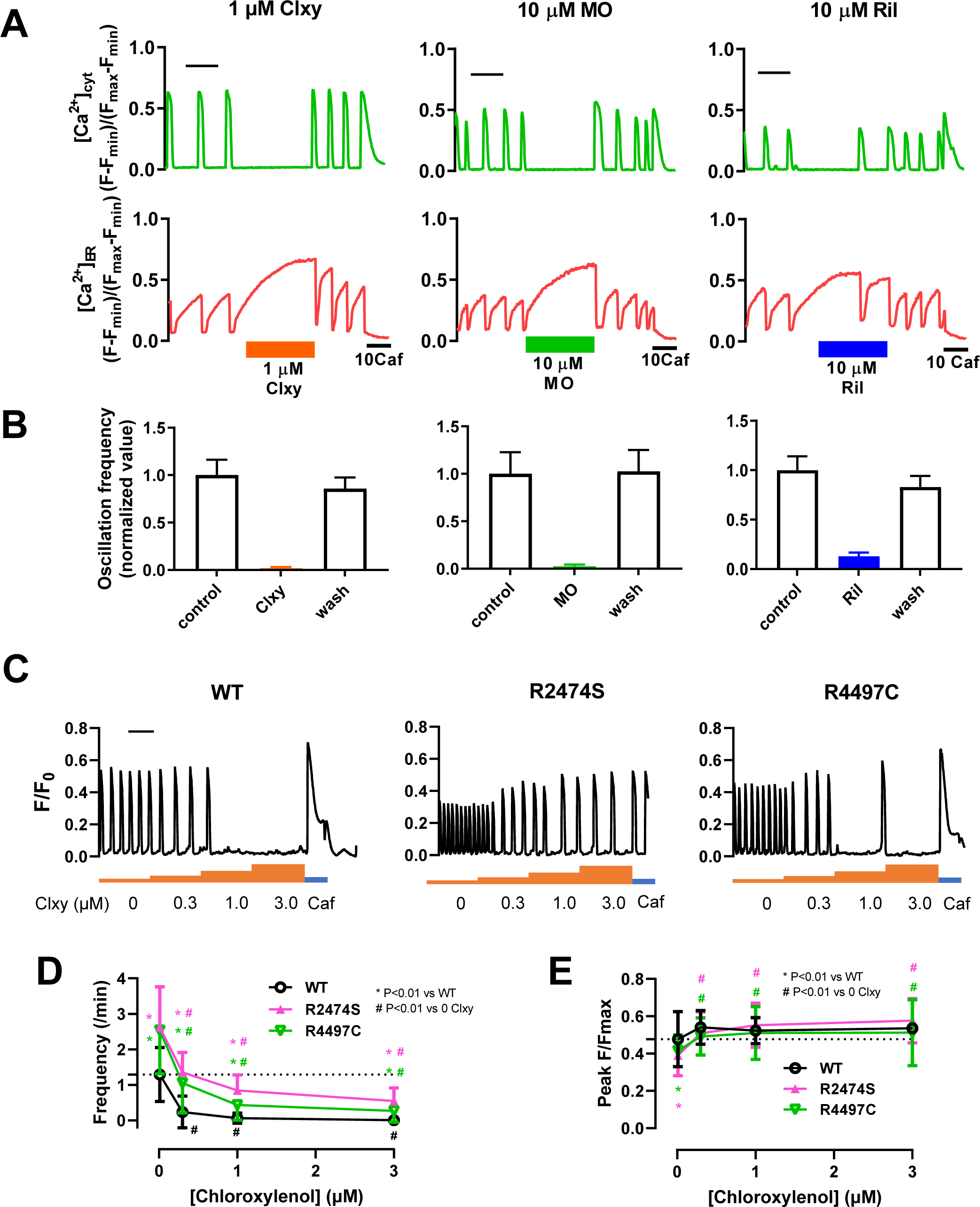
Effects of RyR2 inhibitors on Ca^2+^ oscillations in HEK293 cells expressing WT and mutant RyR2. (A) Representative effects of chloroxylenol (Clxy, left), methyl orsellinate (MO, center), and riluzole (right) on [Ca^2+^]_cyt_ (top) and [Ca^2+^]_ER_ (bottom) signals in single WT RyR2 cells. Drugs were applied for 5 min after 5 min conditioning in normal Krebs solution. Scale bar, 2 min. (B) Effects of chloroxylenol (1 μM, left), methyl orsellinate (10 μM, middle), and riluzole (10 μM, right) on cytoplasmic Ca^2+^ oscillation frequency. The oscillation frequency in individual cells was normalized to the average value before drug treatment (control). Data are mean ± SD (n=24–40 from 2 dishes). (C) Representative dose-dependent effects of chloroxylenol on [Ca^2+^]_cyt_ signals in single WT (left), R2474S (middle), and R4497C (right) cells. Drugs were cumulatively administered for 5 minutes each. Scale bar, 2 min. (D) and (E) Dose-dependent effects of chloroxylenol on Ca^2+^ oscillation frequency (D) and peak [Ca^2+^]_cyt_ transients (E) on WT (black), R2474S (pink) and R4497C (green) cells. Data are mean ± SD (n=67–70 from 2 dishes) and were analyzed by two-way ANOVA with Tukey’s test.

We have previously reported that the enhancement of RyR2 activity by CPVT-linked mutations will result in an increase in Ca^2+^ oscillation frequency and a decrease in the amplitude of Ca^2+^ transients due to the decrease in [Ca^2+^]_ER_ (Kurebayashi et al., 2022). We investigated whether these RyR2 inhibitors could makes the enhanced Ca^2+^ oscillations in CPVT mutant RyR2s close to that of WT RyR2. Figure 4C shows the effects of various concentrations of chloroxylenol, which has the highest affinity for RyR2 among the three compounds, on cytoplasmic Ca^2+^ oscillations in cells expressing WT and CPVT-linked mutant RyR2s. In RyR2-WT cells, Ca^2+^ oscillation was substantially reduced at 0.3 μM and almost disappeared at 1 μM. Compared with the WT cells, mutant RyR2 cells, R2474S and R4497C cells exhibited more frequent with smaller-amplitude Ca^2+^ oscillations in normal Krebs solution (Fig. 4C–E). The addition of chloroxylenol to R2474S and R4497C cells increased the amplitudes of Ca^2+^ transients and decreased the frequencies of Ca^2+^ oscillations in a dose-dependent manner (Fig. 4C–E). Notably, oscillation frequency and peak amplitude of mutant cells in the presence of 0.3 μM chloroxylenol were similar to those of WT cells in the absence of the drug, suggesting that RyR2 inhibitors can ameliorate abnormal Ca^2+^ homeostasis in RyR2-expressing cells at around their IC_50_s.

### Effects of hit compounds on Ca^2+^ signals in isolated cardiomyocytes from disease model mice

The above results demonstrate that the hit compounds suppress RyR2 in HEK293 cells. Next, we investigated whether the hit compounds can suppress abnormal Ca^2+^ signals such as Ca^2+^ waves in cardiomyocytes. For this purpose, we used cardiomyocytes from mice harboring CPVT-linked RyR2 mutation, R420W (Nishio et al., 2006; Okudaira et al., 2014). The R420W cardiomyocytes stimulated at 0.5 Hz showed few Ca^2+^ waves in normal Krebs solution but showed frequent Ca^2+^ waves in the presence of 10^−7^ M isoproterenol (ISO) (Fig. 5A and B, Suppl. video 1). We tested the effects of the RyR2 inhibitors at a concentration near their IC_50_s (i.e., 0.3 μM for chloroxylenol, 3 μM for methyl orsellinate, and 10 μM for riluzole) on ISO-induced Ca^2+^ waves. Figure 5C shows typical Ca^2+^ transients and Ca^2+^ waves in RyR2-R420W cardiomyocytes stimulated at 0.5 Hz. As shown in Fig. 5C-a and D-a, Ca^2+^ waves were observed in normal Krebs and after 5 min of incubation with vehicle (0.1% DMSO). In the presence of 0.3 μM chloroxylenol, cardiomyocytes showed normal Ca^2+^ transient in response to field stimulation; however, the frequency of Ca^2+^ waves dramatically decreased (average 9-fold reduction compared to ISO) (Fig. 5C-b and 5D-b, also see Supple. videos 1 and 2). High resolution videos in the presence of isoproterenol alone and isoproterenol plus chloroxylenol are shown in Suppl. video 3 and Suppl. video 4 respectively. Similar reduction in Ca^2+^ frequencies were also seen in the presence of 3 μM methyl orsellinate (average 10-fold) (Fig. 5C-c and D-c) and 10 μM riluzole (average 14-fold) (Fig. 5C-d and D-d).

**Fig. 5.**
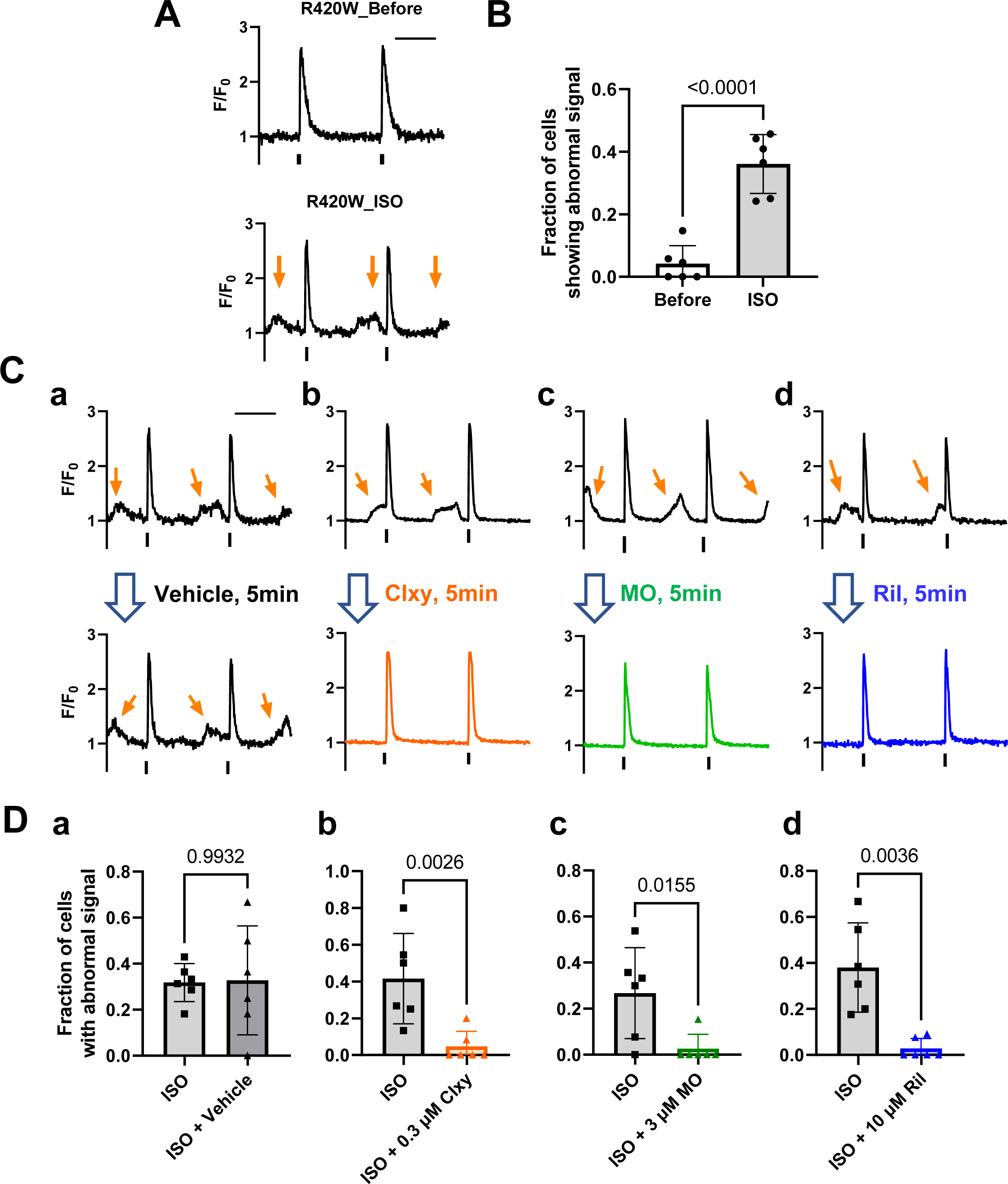
Effects of RyR2 inhibitors on Ca^2+^ signals in cardiomyocytes from RyR2-R420W mice. (A) Representative isoproterenol (ISO)-induced Ca^2+^ signals in R420W mutant cardiomyocytes. Myocytes were field-stimulated at 0.5 Hz. Note that abnormal Ca^2+^ signals, such as diastolic Ca^2+^ waves, were frequently observed in the presence of ISO in R420W cardiomyocytes. Scale bar, 1 sec. (B) Fraction of cells showing abnormal Ca^2+^ signals in R420W cardiomyocytes before and after 5min incubation with ISO. Ca^2+^ signals were measured from 23–58 cells per mouse. Data are mean ± SD (n=6 mice) and were analyzed by unpaired t-test. (C). Representative effects of vehicle (0.1% DMSO) (a), 0.3 μM chloroxylenol (Clxy) (b), 3 μM methyl orsellinate) (MO) (c), and 10 μM riluzole (Ril) (d) on ISO-induced abnormal Ca^2+^ signals in R420W cardiomyocytes. Scale bar, 1 sec. (D) Fraction of cells showing abnormal Ca^2+^ signals in the presence of ISO alone and ISO plus vehicle (0.1% DMSO) (a), 0.3 μM chloroxylenol (Clxy) (b), 3 μM methyl orsellinate (c), and 10 μM riluzole (d). Ca^2+^ signals were measured from 5–15 cells per mouse. Data are mean ± SD (n=6 mice) and were analyzed by unpaired t-test.

We also examined the effects of the three hit compounds on Ca^2+^ waves in cardiomyocytes from dilated cardiomyopathy (DCM) model mice, *tnnt2* ΔK210, which develop progressive severe heart failure and ventricular arrhythmia with aging (Odagiri et al., 2014; Suzuki et al., 2012). These mice exhibit increased phosphorylation levels of RyR2 in the heart, suggesting enhanced RyR2 activity in the *tnnt2* ΔK210 mouse heart (Du et al., 2007). Cardiomyocytes from the DCM mice at 3 months old showed frequent Ca^2+^ waves (19.9±4.6%, n = 6 mice), in normal Krebs solution (Fig. 6A, before drug). Ca^2+^ wave frequencies were measured in normal Krebs solution and then after 5 min of incubation in Krebs solution containing vehicle alone (0.1% DMSO), 0.3 μM chloroxylenol, 3 μM methyl orsellinate, or 10 μM riluzole. None of these compounds inhibited the generation of action-potential-induced Ca^2+^ transients (Fig. 6A). The average Ca^2+^ wave frequency was increased 1.4-fold in control Krebs solution with vehicle (Fig. 6B-a), but the difference was not statistically significant. This increase is likely due to the natural progression of injury over time. Chloroxylenol (Fig. 6A-b and B-b) and riluzole (Fig. 6A-d and B-d) clearly reduced the occurrence of Ca^2+^ waves (average 3.3- and 25-fold compared to the pretreatment, respectively). Methyl orsellinate did not show statistically significant reduction in the proportion of cells showing Ca^2+^ waves (Fig. 6A-c and B-c).

**Fig. 6.**
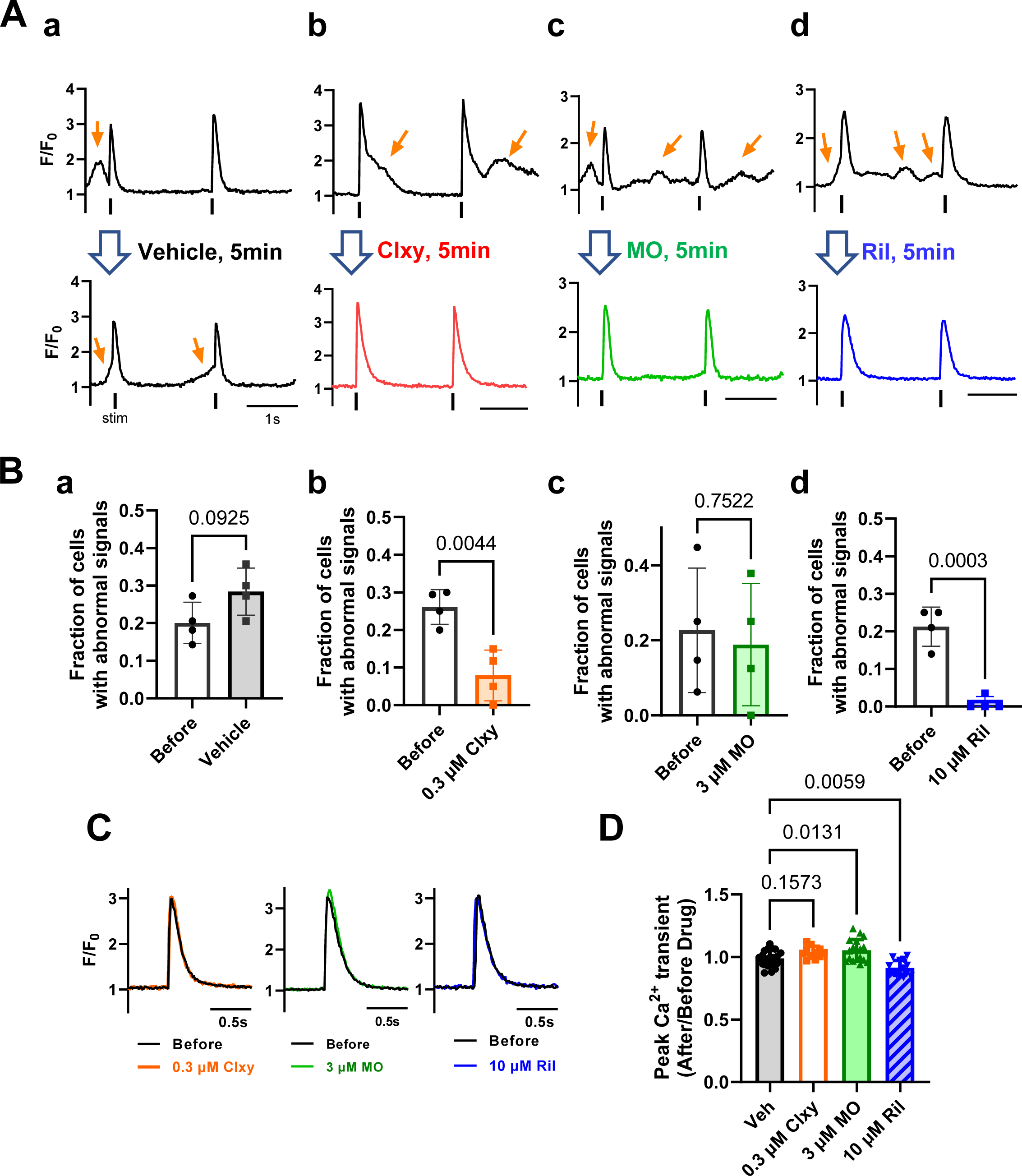
Effects of RyR2 inhibitors on Ca^2+^ signals in cardiomyocytes from *tnnt2* ΔK210 heart failure model mice. (A) Representative abnormal Ca^2+^ signals in *tnnt2* ΔK210 cells. Myocytes were field-stimulated at 0.5 Hz. Note that abnormal Ca^2+^ signals, such as diastolic Ca^2+^ waves, were observed (orange arrows). Scale bar, 1 sec. (B) Fraction of cells showing abnormal Ca^2+^ signals in before and after 5min incubation with (0.1% DMSO) (a), 0.3 μM chloroxylenol (b), 3 μM methyl orsellinate (c), and 10 μM riluzole (d). Data are mean ± SD (n=4 mice, Ca^2+^ signals from 8–30 cells were measured per mouse) and were analyzed by unpaired t-test. (C) and (D) Effects of hit compounds on Ca^2+^ transients in WT cardiomyocytes. (C) Representative traces of Ca^2+^ transients before and after 5 min incubation with 0.3 μM chloroxylenol (a), 3 μM methyl orsellinate (b), and 10 μM riluzole (c). (D) Effects of the three compounds on peak amplitudes of Ca^2+^ transients. Peak amplitudes of Ca^2+^ transients 5 min after application of compounds were normalized to those before the treatments. Data are mean ± SD (n=10–20 from 2 mice) and were analyzed by one-way ANOVA followed by Dunnet’s test.

In Ca^2+^ transient measurements in mutant cardiomyocytes, RyR2 inhibitors suppressed Ca^2+^ waves but did not suppress the generation of action-potential-induced Ca^2+^ transients. Because it is difficult to analyze Ca^2+^ transients in cells that are prone to Ca^2+^ waves, we further examined the effects of these RyR2 inhibitors on Ca^2+^ transients in cardiomyocytes from WT mice. The peak amplitude of Ca^2+^ transients in WT cells was not appreciably altered by these inhibitors. The average amplitudes were only slightly increased by chloroxylenol (mean difference = +4.8%, 95% CI = −1.3 – +11.0%) and methyl orsellinate (mean difference = +6.4%, 95% CI = +1.1 – +11.8%) and decreased by riluzole (mean difference = −7.5%, 95% CI = −1.8 – −13.3%). These results indicate that the RyR2 inhibitors found in this study effectively suppress Ca^2+^ waves at around their EC_50_s without greatly affecting the peak amplitudes and time courses of action-potential-induced Ca^2+^ transients (Fig. 6C and D).

## Discussion

In this study, we found that three compounds—chloroxylenol, methyl orsellinate, and riluzole—have marked RyR2 inhibitory effects, by searching a well-known compound library using ER Ca^2+^ signal-based assay. These compounds reversibly suppressed Ca^2+^ oscillations and elevated [Ca^2+^]_ER_ in RyR2-expressing HEK293 cells and dose-dependently suppressed Ca^2+^-dependent [^3^H]ryanodine binding. Notably, these compounds suppressed Ca^2+^ waves in cardiomyocytes from CPVT model mice and DCM mice. The fact that the three unrelated compounds showed similar effects on RyR2-expressing cells supports the idea that RyR2 inhibitors are promising as antiarrhythmic drugs.

On the basis of the finding that [Ca^2+^]_ER_ is inversely correlated with the channel activity of the RyR1 and RyR2 mutants in HEK293 cells (Kurebayashi et al., 2022; Murayama et al., 2016; Murayama et al., 2015), we recently identified novel RyR1 inhibitors with the screening platform by time-lapse fluorescence measurement of [Ca^2+^]_ER_ using R-CEPIA1er (Murayama et al., 2018). Using the same procedure, we here identified three potent RyR2 inhibitors with IC_50_s of less than 10 μM in the well-known compound library. Because this method targets RyR2 on the ER membrane in living cells, only compounds that can pass through the cell membrane are detected as hits. All the hit compounds identified by our method certainly inhibited [^3^H]ryanodine binding and suppressed abnormal Ca^2+^ waves without affecting action-potential induced Ca^2+^ transients in cardiomyocytes from the disease-linked mouse models. Thus, our method is highly efficient in identifying membrane permeable RyR2 inhibitors needed for clinical use.

Riluzole is a therapeutic agent for amyotrophic lateral sclerosis (ALS), and is known to act on various channels, including neuronal Na^+^ channel (nNaV) and glutamate receptors (Cheah et al., 2010). Although its effect on RyR2 may be one of its non-specific actions, it is characterized by a higher affinity for RyR2 than for RyR1 and RyR3. Interestingly, Radwanski et al. reported that riluzole suppressed diastolic Ca^2+^ release in cardiomyocytes from CPVT linked CASQ2-mutant mice (CASQ2-R33Q) (Radwanski et al., 2015) and suggested that nNav contributes to arrhythmogenic diastolic Ca^2+^ release, thereby providing the mechanism behind its potential for antiarrhythmic therapy. Since the 10 µM riluzole they used has a strong RyR2 inhibitory effect, at least part of the riluzole’s effect on Ca^2+^ abnormalities in cardiomyocyte may be due to the RyR2 inhibitory action. Because therapeutic plasma concentration is reported to be approximately 1∼2 µM (Sarkar et al., 2017), slightly higher doses of the drug may be required for effective suppression of arrhythmia.

Chloroxylenol is an antiseptic and disinfecting agent, and is most effective against gram-positive bacteria. The chemical structure of chloroxylenol is similar to those of 4-CMC and 4-CEP, well-known activators of RyR1, which stimulates Ca^2+^-dependent [^3^H]ryanodine binding with an EC_50_ of approximately 100 μM or higher (Westerblad et al., 1998; Zorzato et al., 1993). Chloroxylenol was found to suppress RyR2 with an IC_50_ of 0.3 μM, but it showed a trend of activation at a concentration of 30 μM or more, whereas it only activated RyR1 without any inhibitory effect. Because this compound distinguishes between RyR1 and RyR2 at low concentrations, it would be interesting to know the binding sites at low and high concentrations in RyR2 molecules. Methyl orsellinate also showed dose-dependent biphasic effect similar to that of chloroxylenol (Fig. 2). Chloroxylenol and methyl orsellinate may act on the same binding site in RyR2.

Notably, all of these compounds suppressed Ca^2+^ waves in cardiomyocytes from RyR2 R420W and *tnnt2* ΔK210 mice at around their IC_50_s, but only slightly affected peak amplitudes of Ca^2+^ transients in cardiomyocytes from WT mice. This important property can be explained by the mechanism of inhibition of the compounds (Fig. 3): they all shift Ca^2+^ dependence to the right and mildly suppress the maximal activity, leading to a greater suppression of Ca^2+^ release at lower [Ca^2+^]_cyt_ than at higher [Ca^2+^]_cyt_. As a result, local Ca^2+^ waves/sparks generated at low [Ca^2+^]_cyt_ may be effectively suppressed, but action-potential-evoked Ca^2+^ transients caused by massive Ca^2+^ influx via LTCC may be less suppressed. Interestingly, these compounds were able to ameliorate the frequent Ca^2+^ release by gain-of-function mutant RyR2 to a level closer to that of the WT in HEK293 cells.

It is widely believed that drugs that suppress RyR2 have an antiarrhythmic effect, and indeed several drugs have been reported to modify RyR2 activity (Szentandrassy et al., 2022). However, in many cases, these drugs also have strong effects on other target proteins, and so this concept has yet to be definitively confirmed. Flecainide is an effective antiarrhythmic drug that mainly targets Na^+^ channels, but its action on RyR2 has been reported to contribute to its antiarrhythmic effect (Bannister et al., 2022; Salvage et al., 2022; Watanabe et al., 2009). A β-blocker, carvedilol, has also been reported to suppress abnormal Ca^2+^ release via RyR2 (Zhou et al., 2011). There is thus a need to clarify whether and how RyR2 inhibitors exhibit antiarrhythmic effects using strong RyR2 inhibitory effects.

EL20, which is an analog of tetracaine, has been reported to potently inhibit abnormal Ca^2+^ release from RyR2 mutant mice and induced pluripotent stem cell derived cardiomyocytes (iPSC-CM) (Klipp et al., 2018; Word et al., 2021). This compound strongly inhibited single channel activity of RyR2 (IC_50_=8.2 nM) in the absence of calmodulin. However, in the presence of calmodulin, EL20 showed a much weaker effect (IC_50_=3–10 μM) on the single channels and on Ca^2+^-dependent [^3^H]ryanodine binding in cardiac microsomes. The unnatural verticilide exhibits antiarrhythmic activity and appears to be RyR2 specific, but its inhibitory effect on [^3^H]ryanodine binding is partial (Batiste et al., 2019). A group of compounds such as Rycal drugs that stabilize the association of FK binding protein 12.6 (FKBP12.6/calstabin2) with RyR2 has also been reported to ameliorate abnormal Ca^2+^ release in cardiomyocytes (Andersson and Marks, 2010). Compared with these drugs/compounds previously reported to suppress RyR2 activity, the compounds found in our study clearly suppress [^3^H]ryanodine binding, especially at Ca^2+^ concentrations of less than 10 μM. Importantly, the inhibitory effects of the three compounds were not mediated by associated proteins (FKBP12.6 and calmodulin) or phosphorylations. They inhibited FKBP12.6-unbound RyR2 in HEK293 cells that do not express FKBP12.6. They also appear to suppress both calmodulin-unbound and calmodulin-bound RyR2, as the amount of calmodulin in RyR2 expressing HEK293 cells is insufficient in molar ratio compared to the amount of RyR2 (Gao et al., 2023). Furthermore, they appear to inhibit phosphorylated and non-phosphorylated RyR2s in a similar manner, as they suppressed both phospho-null (RyR2-S3A) and phospho-mimetic mutants (RyR2-S3D) (Supplemental Fig. 1).

In addition to their use as antiarrhythmic drugs, RyR2 selective inhibitors are expected to be beneficial agents for the other diseases. In the hearts of HF and diabetic patients, various ion channels and regulatory proteins have been reported to be modulated. Among them, RyR2 can be modulated by association/dissociation of FKBP12.6 (calstabin2), calmodulin, triadin/calsequestrin, calcineurin, and transmembrane protein 38A (TMEM-38A) (Kansakar et al., 2021; Santulli et al., 2018; Yuan et al., 2014), as well as phosphorylation, oxidation, S-nitrosylation and glycation of RyR2 (Gambardella et al., 2022; Kansakar et al., 2021; Kobayashi et al., 2021; Tian et al., 2021). Furthermore, RyR2 have been also implicated in the diseases in brain such as Alzheimer’s disease and epilepsy (Lacampagne et al., 2017; Lehnart et al., 2008; Lieve et al., 2019) and in pancreas (Santulli et al., 2018). RyR2-selective inhibitors might help to treat these diseases.

### Limitations and perspectives

Chloroxylenol and methyl orsellinate have activating effects on RyR1, and riluzole has insufficient affinity for RyR2 inhibition, and thus we are not ready to test these compounds as antiarrhythmic agents on animals in their current form. In this regard, it is necessary to develop compounds with higher affinity and selectivity for RyR2 by structural modification of the three lead compounds. For that purpose, analysis of ultrafine structure of compound-bound RyR2 by cryo electron microscopy would be helpful (Iyer et al., 2020; Kobayashi et al., 2022; Santulli et al., 2018; Woll and Van Petegem, 2022). Our screening method should greatly contribute to the search for novel RyR2 inhibitors as a new category of antiarrhythmic drugs.

## Acknowledgments

The authors are grateful to Dr Montserrat Samso for the suggestion to create the RyR2-S3A and RyR2-S3D cell lines. The authors thank Ikue Hiraga, and Mirei Takahashi for their technical assistance and Dr. Yutaka Hirata for helpful advice on usage of the mutant mice. We are grateful for the Center for Biomedical Research Resources and Laboratory of Radioisotope Research, Research Support Center, Juntendo University Graduate School of Medicine. The authors also thank Edanz (https://jp.edanz.com/ac) for editing a draft of this manuscript.

## Data availability

All data supporting the findings in this study are available upon request.

## Authorship contributions

M. Takenaka, M. Kodama, T. Murayama contributed equally to this work. Participated in research design: T. Murayama, T. Sakurai, H. Kagechika and N. Kurebayashi; Conducted experiments: M. Takenaka, M. Kodama, T. Murayama and N. Kurebayashi; Contributed new reagents or analytic tools: M. Ishigami-Yuasa, S. Mori, R. Ishida, H. Kagechika, J. Suzuki, K. Kanemaru, M. Iino A. Miura, H. Nishio and S. Morimoto; Performed data analysis: M. Takenaka, M. Kodama, T. Murayama, M. Ishigami-Yuasa, M. Sugihara and N. Kurebayashi analyzed the experimental data; Wrote or contributed to the writing of the manuscript: M. Takenaka, M. Kodama, T Murayama, and N. Kurebayashi. All authors discussed the results and approved the final version of the manuscript.

## Abbreviations

ARVD: arrhythmogenic right ventricular dysplasia
CPVT: catecholaminergic polymorphic ventricular tachycardia
[Ca^2+^]_ER_: ER Ca^2+^ concentrations
[Ca^2+^]_cyt_: cytosolic Ca^2+^ concentrations
4-CEP: 4-chloro-3-ethylphenol
4-CMC: 4-chloro-m-cresol
DCM: dilated cardiomyopathy
DMSO: dimethyl sulfoxide
IVF: idiopathic ventricular tachycardia
LQTS: Long QT syndrome
LTCC: L-type Ca^2+^ channel
NCX: Na^+^-Ca^2+^ exchanger
RyR: Ryanodine receptor
95% CI: 95% Confidence interval

## Footnotes

This work was supported by JSPS KAKENHI Grant Number 19K07105 and 22K06652 to N.K., 19H03404 and 22H02805 to T.M., 22K15244 to R.I., the Practical Research Project for Rare/Intractable Diseases (19ek0109202 to N.K.) from the Japan Agency for Medical Research and Development (AMED), Platform Project for Supporting Drug Discovery and Life Science Research (Basis for Supporting Innovative Drug Discovery and Life Science Research (BINDS) (JP19am0101080 to T.M. and N.K. and JP20am0101098 to H.K.), the Cooperative Research Project of Research Center for Biomedical Engineering (to H.K.), an Intramural Research Grant (2-5 to T.M.) for Neurological and Psychiatric Disorders from the National Center of Neurology and Psychiatry to T.M., and the Vehicle Racing Commemorative Foundation (6303 to T.M.).

The authors declare no competing financial interests.

