## Supplemental Fig 1 for "Screening for novel RyR2 inhibitors by ER Ca^2+^ monitoring"

MOLPHARM-AR-2023-000720

Supplemental Data
Molecular Pharmacology

**Screening for novel RyR2 inhibitors by ER $Ca^{2+}$ monitoring**

Mai Takenaka*, Masami Kodama*, Takashi Murayama*, Mari Ishigami-Yuasa, Shuichi Mori, Ryosuke Ishida, Junji Suzuki, Kazunori Kanemaru, Masami Sugihara, Masamitsu Iino, Aya Miura, Hajime Nishio, Sachio Morimoto, Hiroyuki Kagechika[†], Takashi Sakurai and Nagomi Kurebayashi[†]

* indicates co-first authors

† indicates corresponding authors

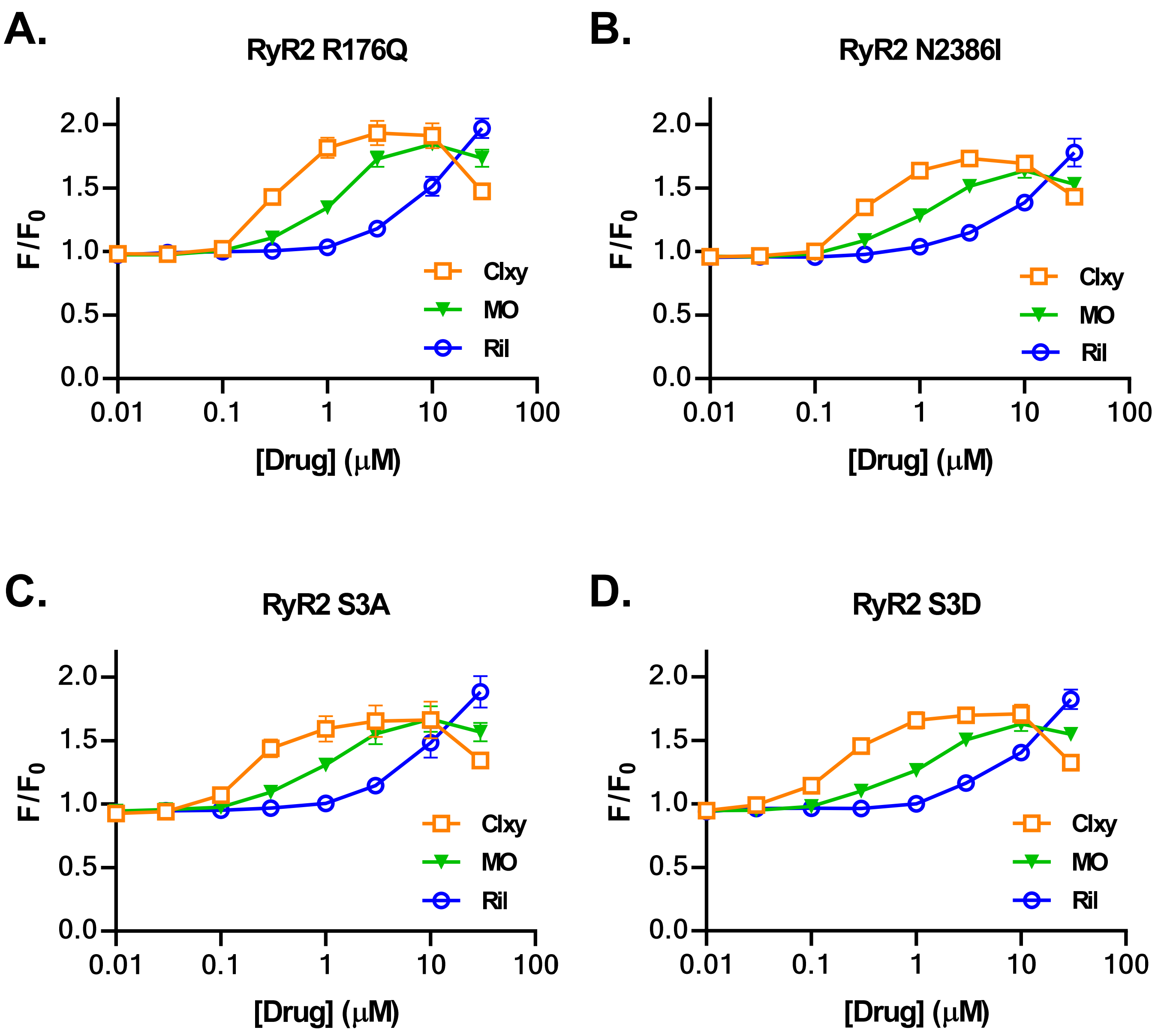


**Fig. S1 Dose-dependent effects of hit compounds on R-CEPIA1er signals in mutant RyR2-expressing cells determined with FlexStation3.** (A) RyR2-R176Q, (B) RyR2-N2386I, (C). RyR2 phospho-null triple mutant S2807A/S2813A/S2030A (RYR2 S3A), (D) RyR2 phospho-mimetic triple mutant of RyR2 S2807D/S2813D/S2030D (RyR2 S3D). Data are mean ± SD (n=3–5).
